# Pan-cancer benchmarking reveals complementary copy number signatures with distinct multi-omic predictability

**DOI:** 10.64898/2026.07.20.739333

**Authors:** Marco Rota Negroni, Ilaria Billato, Chiara Romualdi

## Abstract

Copy number signatures provide compact representations of the processes that shape cancer genomes, but signatures derived with different feature encodings are often interpreted as if they were interchangeable. We established a matched-sample pan-cancer benchmark of three major copy number signature compendia, comparing their activity structure, cross-framework concordance, patient stratification, outcome associations, and predictability from non-copy-number molecular data. Signature-level concordance was sparse and concentrated in a limited set of biologically related patterns. Clustering of high-activity signatures produced distinct patient partitions with limited overlap between compendia, although one cluster in each framework showed a directionally favorable outcome association after accounting for cancer-type-specific baseline hazards. Prediction from gene expression, DNA methylation, somatic mutations, age, and tumor purity was strongly framework dependent: test-set F1 scores were 0.93 for Drews, 0.80 for Steele, and 0.24 for Tao. Gene expression provided the largest contribution and largely retained the performance of the full models. These results show that compendium choice is an analytical decision rather than an interchangeable preprocessing step. The benchmark provides a reproducible framework for selecting and interpreting copy number signature representations in pan-cancer studies.

## Introduction

Somatic copy number alterations range from focal gains and deletions to chromosome-arm changes and whole-genome doubling. Together, these events reshape gene dosage, disrupt tumor suppressors, amplify oncogenes, and record the evolutionary processes acting on a cancer genome (1). Copy number signatures summarize recurrent combinations of these alterations and can therefore provide a more interpretable representation of genome instability than individual copy number events. Initial work in ovarian cancer demonstrated that such signatures can recover distinct mutational processes and clinically relevant genome states (2).

Three pan-cancer studies subsequently produced broad copy number signature compendia using different feature encodings and analytical choices. Drews et al. described 17 chromosomal instability (CIN) signatures based on breakpoint density, segment size, copy number change, and oscillating patterns, with proposed links to mitotic errors, homologous recombination deficiency (HRD), replication stress, and other instability processes (3). Steele et al. derived 21 signatures from allele-specific copy number profiles encoded by total copy number, loss of heterozygosity (LOH), and segment size, capturing states that include diploidy, whole-genome doubling, aneuploidy, focal and chromosome-scale LOH, chromothripsis, and HRD (4). Tao et al. defined 20 signatures using absolute copy number, LOH, segment size, and local copy number context, providing a granular description of copy number morphology across whole-genome sequencing and single-nucleotide polymorphism array data (5). Clinical investigations have further shown that selected CIN signatures can predict chemotherapy resistance in specific settings (6).

These compendia address related biological questions but do not encode the same objects. A signature in one framework may represent a discrete instability mechanism, whereas a signature in another may summarize ploidy, LOH, or a composite morphology. Consequently, apparent similarity between signatures can reflect shared biological aetiology, overall alteration burden, or concordant inactivity rather than mathematical equivalence. Conversely, signatures that are weakly correlated at the individual level may still generate patient groups with comparable biological or clinical properties. Recent extensions of copy number signature analysis to targeted clinical panels (7) and attention-based learning of cancer-associated copy number patterns (8) make the comparability of signature representations an increasingly important methodological question.

A matched-sample benchmark is therefore needed before copy number signatures can be combined across studies or selected for downstream analyses. Such a benchmark should evaluate several levels of agreement: the activity distributions generated by each framework; signature-level concordance in the same tumors; the patient partitions induced by high-activity signatures; the persistence of outcome associations after accounting for cancer type; and the extent to which each representation can be recovered from other molecular measurements. Predictability from gene expression, DNA methylation, and somatic mutation profiles is particularly informative in this context. Pan-cancer analyses have shown that somatic copy number alterations are major contributors to gene-expression variation, providing a biological basis for testing cross-modal predictability (9). It tests whether a signature-derived state has a reproducible molecular correlate, while avoiding the stronger and unsupported assumption that these data can replace direct copy number profiling.

Here, we compare the Drews, Steele, and Tao compendia across The Cancer Genome Atlas (TCGA). The study addresses three questions: how concordant and interchangeable are the three signature representations in matched tumors; whether their cluster-level patient stratifications preserve comparable biological and clinical information; and whether those framework-defined states differ in their multi-omic predictability. By integrating matched-sample correlations, binary agreement, model-based clustering, cancer-type-stratified survival analysis, and held-out machine learning evaluation, we show that the three compendia are complementary rather than interchangeable and that their downstream predictability is itself framework dependent (Figure 1).

**Figure 1.**
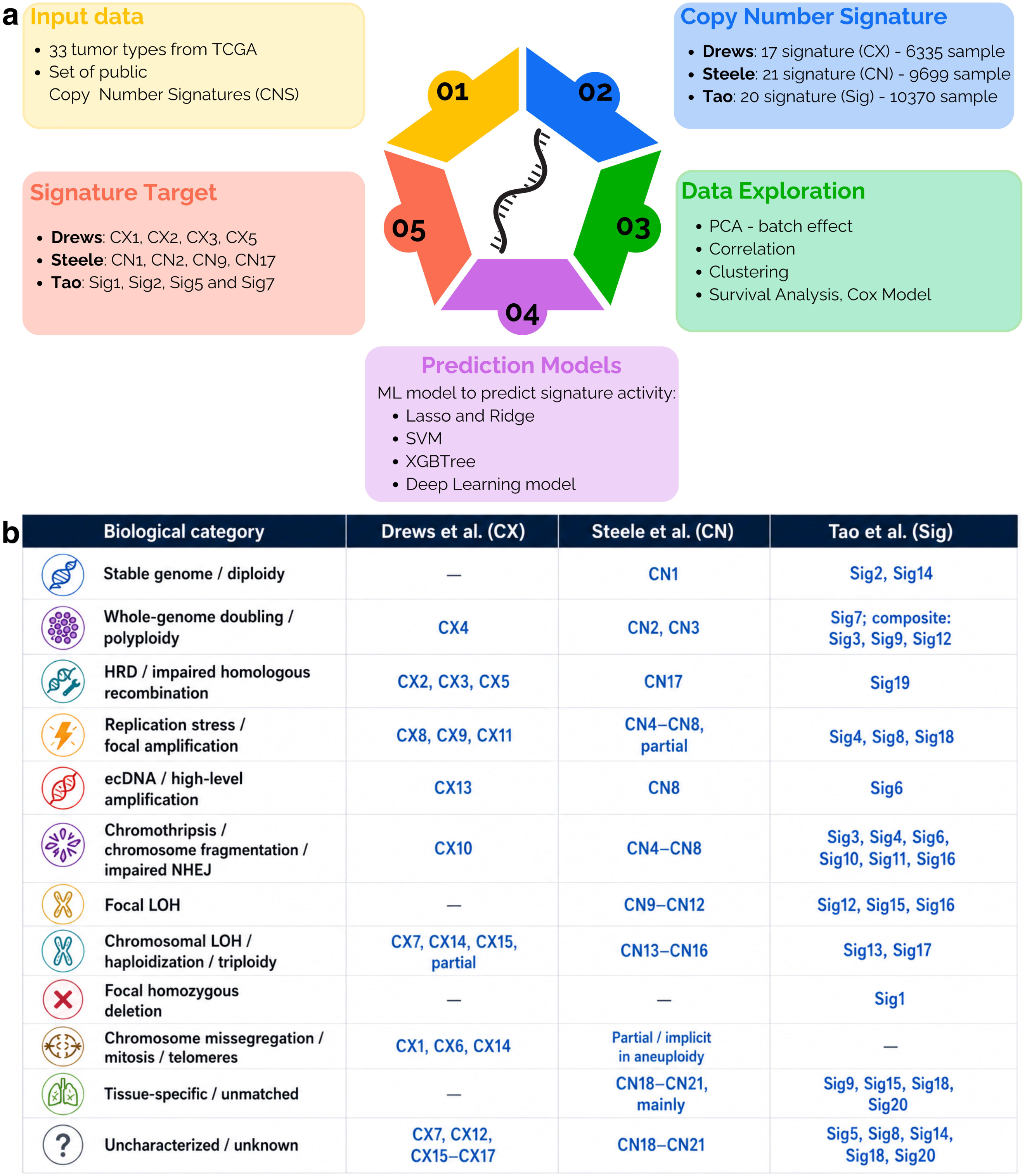
Study design and qualitative comparison of copy number signature frameworks. (a) Workflow showing the three source compendia, selection of high-activity signatures, exploratory and concordance analyses, patient clustering, cancer- type-stratified survival analysis, and multi-omic prediction. The signature comparison, clustering, and survival analyses covered 33 TCGA disease cohorts, including LAML. Predictive modeling was restricted to 32 primary solid-tumor cohorts (TCGA sample type 01), thereby excluding LAML. (b) Qualitative mapping of signatures to proposed biological or morphological categories. Assignments summarize the interpretations in the source studies and do not imply mathematical equivalence or mutually exclusive aetiology.

## Materials and Methods

### Study design and data sources

Precomputed copy number signature activity matrices were obtained from the repositories accompanying the original Drews, Steele, and Tao publications (3–5). The matrices contained 17 Drews signatures for 6,335 tumors, 21 Steele signatures for 9,699 tumors, and 20 Tao signatures for 10,370 tumors. Each compendium covered 33 cancer types from TCGA; sample availability by cancer type is reported in Supplementary Table 1. Signature activities are represented on a relative scale from 0 to 1.

Throughout the text, the three compendia are referred to by the first author’s name. Original signature identifiers were retained: CX for Drews, CN for Steele, and Sig for Tao. Analyses of within-compendium distributions, cross-compendium concordance, patient clustering, and survival retained acute myeloid leukemia (LAML) and therefore covered all 33 TCGA disease cohorts represented in the signature matrices. TCGA patient identifiers were normalized to 12-character barcodes before matching. Cross-compendium analyses were restricted to the 5,881 tumors represented in all three compendia.

Gene expression and DNA methylation data were obtained through the *curatedTCGAData* Bioconductor resource (10). Somatic mutations were obtained from the Multi-Center Mutation Calling in Multiple Cancers (MC3) public mutation annotation file described by Hoadley et al. (11). Harmonized overall survival (OS), progression-free interval (PFI), age, and other clinical variables were obtained from the pan-cancer clinical resource of Liu et al. (12). Tissue source site annotations were obtained from the Genomic Data Commons code tables. All analyses used de-identified, publicly available data; ethical approval and informed consent were obtained by the original contributing studies, and no new participants were recruited for this secondary analysis.

### Signature activity architecture and sources of variation

Signature activity distributions were examined across all samples in each compendium. Principal component analysis (PCA) was applied to the signature matrices without feature scaling to identify signatures that contributed most strongly to the dominant axes of variation. The four signatures with consistently prominent activity and variance contributions in each compendium were retained for cluster-level analyses: CX1, CX2, CX3, and CX5 for Drews; CN1, CN2, CN9, and CN17 for Steele; and Sig1, Sig2, Sig5, and Sig7 for Tao.

Potential technical and biological sources of variation were assessed using tissue source site and cancer type. For these analyses, signature values were scaled before PCA. Separate linear models were fitted to each of the first five principal components, with either tissue source site or cancer type as the explanatory variable. The resulting coefficients of determination were used as descriptive measures of the variance associated with each factor.

### Cross-compendium concordance benchmark

Within- and cross-compendium concordance was first quantified using Pearson correlation coefficients computed over matched samples. Because many signatures had zero-inflated or strongly skewed activity distributions, a complementary binary comparison was performed. For each signature, values were classified as active when they were greater than zero if the median was zero, or greater than or equal to the median otherwise.

Because both activity and inactivity define informative signature states, each signature was represented as the set of patient-state assignments generated by its binary classification. Similarity between two signatures was quantified as the Jaccard similarity of these assignments, A/(2N - A), where A is the number of patients assigned the same state (active-active or inactive-inactive) and N is the number of matched patients. Thus, concordant active and concordant inactive classifications contributed symmetrically to agreement. This patient-state Jaccard measures concordance of binary patient classification and does not by itself establish coactivation or mechanistic equivalence. Agreement matrices were computed across all signatures and across the subset of high-activity signatures.

### Model-based clustering of signature activities

Patient groups were identified separately within each compendium using the four high-activity signatures defined above. Exploratory k-means solutions across increasing values of k were examined using clustering trees to visualize the movement of samples between resolutions and guide the candidate cluster range (13; Supplementary Figure 2). Final classifications were obtained by Gaussian finite mixture modeling with the *mclust* framework (14), considering two to four mixture components and selecting the number of components and covariance parameterization by the Bayesian information criterion. The retained solutions comprised four clusters for Drews and Steele and three clusters for Tao. Cluster labels are compendium specific and do not imply equivalence between identically numbered clusters.

Tumor-type composition was summarized as the relative frequency of each cluster within each cancer type. Cross-compendium overlap was evaluated after matching patient identifiers, both across all clusters and specifically for Clusters 1 and 2, which showed the most divergent outcome profiles.

### Survival analyses

Kaplan-Meier curves were used to describe OS and PFI across signature-derived clusters. To test whether cluster associations persisted beyond differences in cancer-type composition, separate Cox proportional hazards models were fitted for each compendium and endpoint:

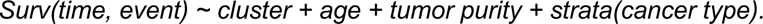

Cancer type was used as a stratification variable, allowing each cancer type to have its own baseline hazard while estimating a common cluster effect. Cluster 1 was the reference group, and age and tumor purity were included as continuous covariates. The complete-case sample sizes were 6,111 tumors with 2,298 events for Drews OS, 6,092 tumors with 2,629 events for Drews PFI, 9,480 tumors with 3,181 events for Steele OS, 9,370 tumors with 3,532 events for Steele PFI, 9,052 tumors with 2,952 events for Tao OS, and 8,945 tumors with 3,229 events for Tao PFI.

Hazard ratios (HRs) are reported with 95% confidence intervals (CIs). Two-sided Wald P values for the 16 cluster coefficients estimated across the six compendium-endpoint models were adjusted as a single family using Holm’s method, which controls the family-wise error rate. Effect sizes and CIs were emphasized when comparing frameworks.

### Multi-omic data preprocessing

For predictive modeling, molecular samples were restricted to TCGA sample-type code 01 (Primary Solid Tumor). LAML samples, which are represented as primary blood-derived cancer rather than sample type 01, were therefore not included in the predictive cohort. Samples were then retained when both gene expression and DNA methylation measurements were available. Low-variance features in the bottom 20% of the coefficient-of-variation distribution were removed, followed by removal of highly correlated features with an absolute Pearson correlation greater than 0.90. Age and tumor purity were included as additional numeric variables. Cancer type was retained only for stratified data splitting and descriptive evaluation and was not included as a model predictor.

Silent somatic mutations were excluded. Mutation dimensionality was reduced following the pathway-based aggregation strategy described by Becchi et al. (15), using KEGG pathway annotations (16); genes without pathway annotations were retained as individual variables. The final predictor space contained more than 35,000 variables: 12,653 gene expression features, 9,227 DNA methylation features, 13,340 gene-level mutation features, 272 pathway-level mutation features, age, and tumor purity.

### Methylation harmonization

TCGA methylation samples were generated using either the 27k or 450k Illumina array, and the 450k platform additionally contains Infinium I and II probe designs with different dynamic ranges (17,18). Data from the two platforms were normalized separately. The 27k probes were mapped to corresponding 450k positions, probes containing common single-nucleotide polymorphisms were removed, and a shared probe set was retained.

The minfi package (19) and beta-mixture quantile normalization (20) were used to harmonize probe-type distributions. The effect of normalization was assessed in acute myeloid leukemia, the only disease cohort for which the same samples were represented on both platforms, using PCA projections, beta-value distributions, and standard-deviation profiles (Supplementary Figure 6). LAML was used solely for this technical cross-platform assessment and was not included in the harmonized methylation matrix used for predictive modeling.

### Predictive modeling benchmark

Two predictive tasks were evaluated. First, continuous activities of the four high-activity signatures in each compendium were modeled using Lasso and Ridge regression, support vector machines with linear and radial kernels, extreme gradient boosting, and a fully connected deep learning model implemented with *Flexynesis* (21). Performance was quantified using mean squared error, mean absolute error, and the coefficient of determination.

Second, the primary comparative task modeled membership in Cluster 1 versus Cluster 2 using extreme gradient boosting. These clusters were selected because they displayed the most divergent outcome profiles within each compendium. Samples were divided into 80% training and 20% held-out test sets using cancer-type-stratified sampling and a fixed random seed of 1234. Cancer type was used only for this stratification and was not included in the predictor matrices. Hyperparameters were selected by 10-fold cross-validation in the training set. Cluster 1 was treated as the positive class, and its test-set F1 score was used as the primary classification metric because the cluster classes were imbalanced. Accuracy was also summarized by cancer type for descriptive purposes when at least four test samples were available but was not used as the principal cross-compendium metric because it can be inflated by class imbalance.

Feature importance was quantified by the gain contributed by each variable across the fitted trees. Variables were ranked, and the smallest set accounting for at least 80% of cumulative gain was retained for interpretation. Reactome over-representation analysis was applied to gene-based features, with false-discovery-rate-adjusted P values below 0.05 considered significant. The full multi-omic benchmark was complemented by models restricted to gene expression features to test how much predictive information was retained in the most widely available molecular layer.

### Reproducibility and statistical software

Analyses were conducted in R and Python. PCA and linear models used the *prcomp* and *lm* functions in R. Correlations used the *cor* function. Survival analyses used the *survival* and *survminer* packages. Machine learning models used *caret*, *xgboost*, and *Flexynesis*. Model-based clustering used *mclust*. Analysis code, processed modeling inputs, and documented notebooks are publicly available as described in the Data Availability section.

## Results

### Benchmark design reveals different activity architectures across compendia

The benchmark comprised 58 signatures across three published compendia and 33 cancer types (Figure 1a, b). The available sample sets differed substantially: 6,335 tumors for Drews, 9,699 for Steele, and 10,370 for Tao (Supplementary Table 1). We therefore separated analyses that required matched tumors from analyses designed to characterize each compendium at its full available scale. No compendium was treated as a gold standard, because each was generated from a different copy number encoding and was designed to resolve different aspects of genome instability.

Most signatures had low or zero activity in most tumors (Figure 2a-c). Four signatures dominated the Drews activity landscape (CX1, CX2, CX3, and CX5), and four dominated Steele (CN1, CN2, CN9, and CN17). Tao activities were more evenly distributed, although Sig1, Sig2, Sig5, and Sig7 showed the strongest overall contributions. Principal component loadings recapitulated these differences (Figure 2d-f): variance in Drews and Steele was concentrated in a small number of signatures, whereas Tao showed a broader contribution from multiple signatures.

**Figure 2.**
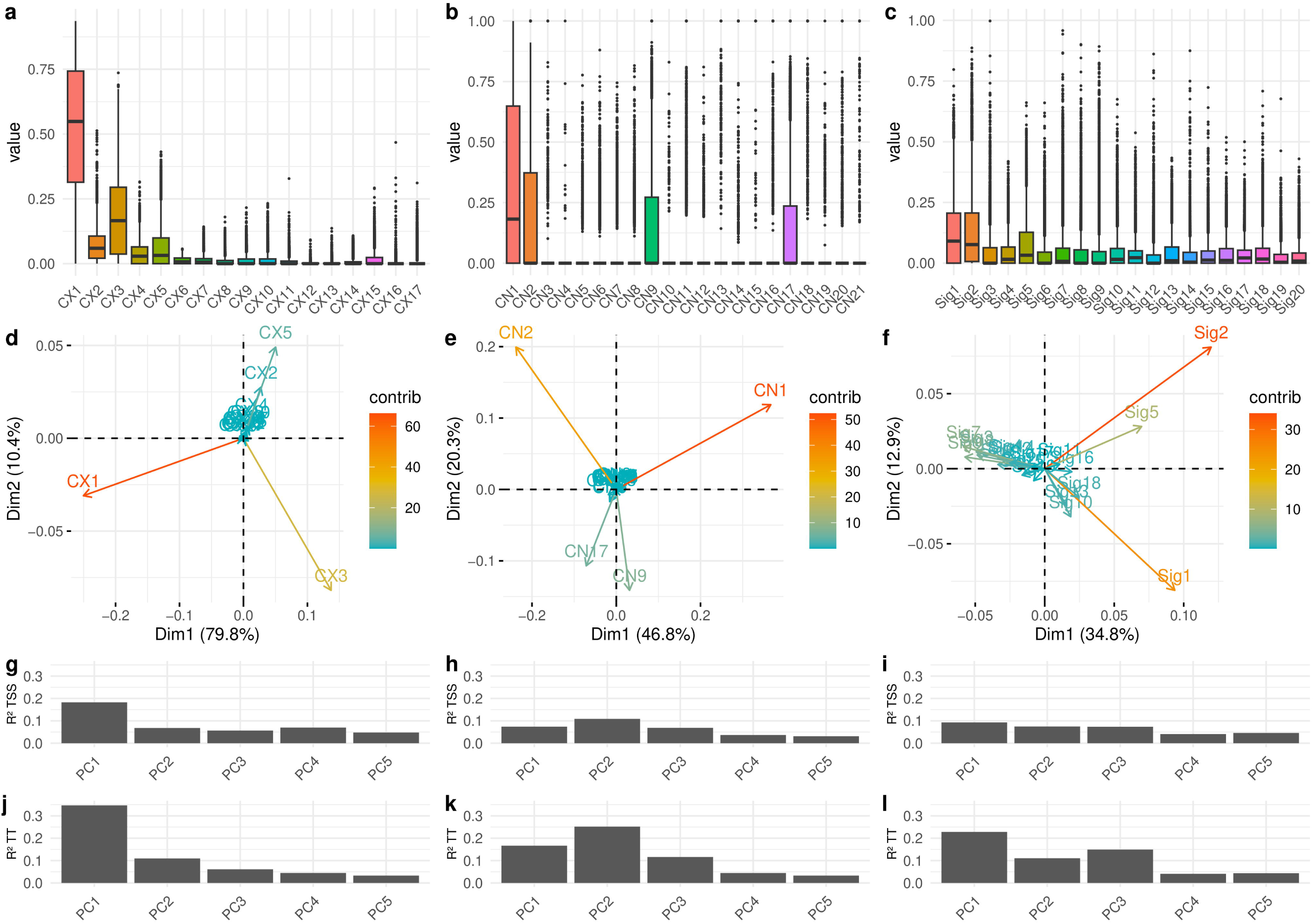
Activity distributions and sources of variation across the three compendia. (a-c) Activity distributions for the Drews, Steele, and Tao signatures. (d-f) Contributions of signatures to the first two principal components. (g-i) Proportion of variance in each of the first five principal components explained by tissue source site in separate linear models. (j-l) Corresponding proportion of variance explained by cancer type. The first five components explain more than 90% of variance in Drews and Steele and more than 70% in Tao.

Because TCGA samples were collected across multiple tissue source sites, we examined whether the leading axes of signature variation were associated more strongly with collection site or with cancer type. Separate linear models were fitted to each of the first five principal components using either tissue source site or cancer type as the explanatory variable. Cancer type accounted for up to approximately 30% of the variance, with the strongest association observed in the Drews compendium (Figure 2j-l). Tissue source site accounted for less variance, reaching approximately 15% for the first principal component and remaining below 10% for the subsequent components (Figure 2g-i). These results indicate that the dominant signature patterns were more closely associated with cancer type than with collection site. Because the two factors were evaluated in separate models, however, their effects cannot be fully distinguished. The observed cancer-type structure motivated cancer-type stratification in the downstream survival and predictive analyses.

### Signature-level concordance is selective and framework dependent

Within each compendium, correlations reflected the genome states emphasized by that framework rather than showing broad coactivation across signatures (Figure 3a,c,e). In the Drews compendium, the strongest negative correlations involved CX1, particularly with the HRD-associated signatures CX2, CX3, and CX5. In Steele, the strongest contrast was between CN1 and CN2 (r = -0.50), consistent with their representation of predominantly diploid and tetraploid genome states. Tao showed a more distributed correlation structure, with negative relationships among Sig1, Sig2, Sig3, and Sig5 and a positive association between Sig2 and Sig5 (r = 0.53), in line with its more granular representation of copy number morphology.

**Figure 3.**
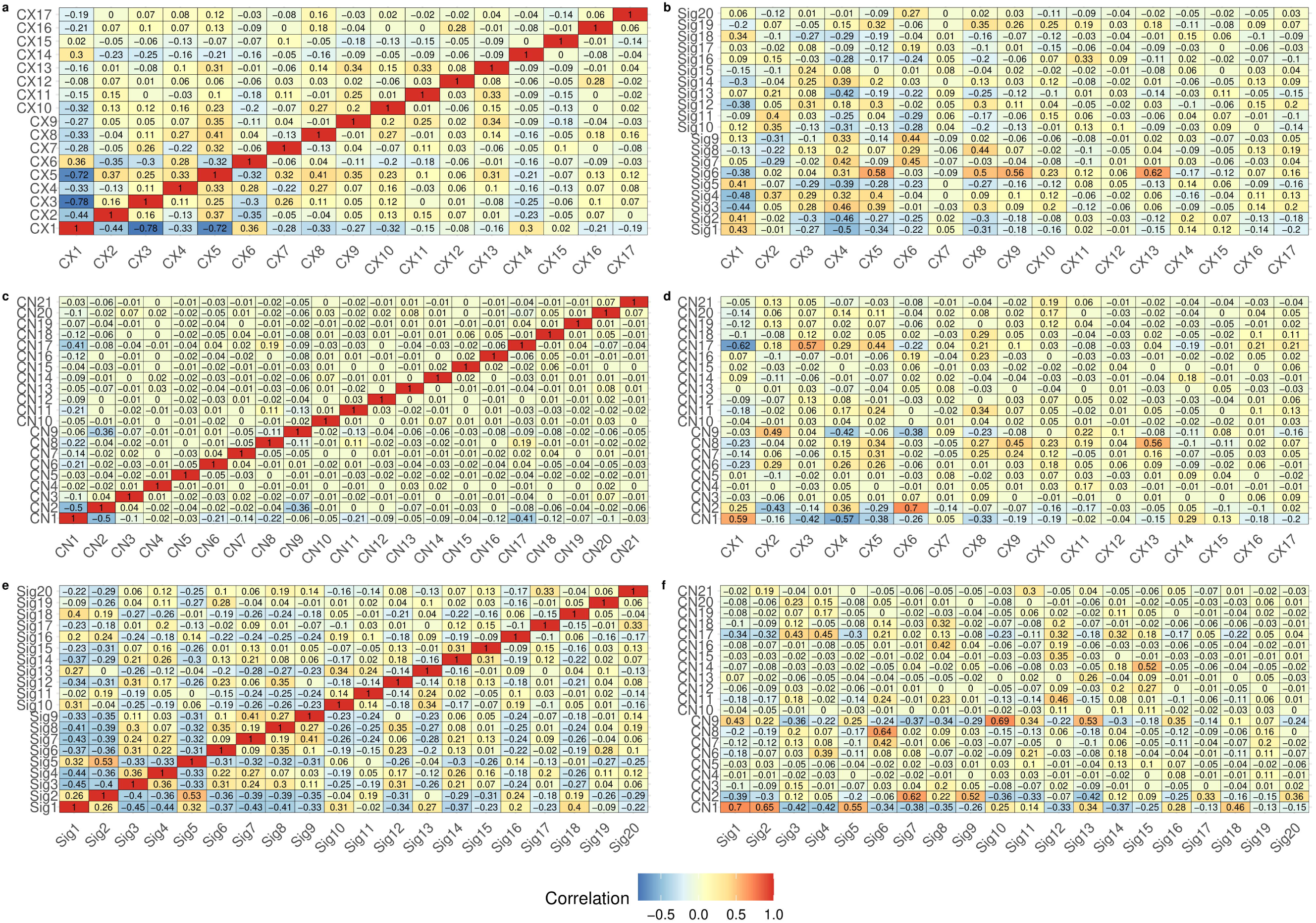
Pearson correlation benchmark within and across copy number signature compendia. (a,c,e) Within- compendium correlation matrices for Drews, Steele, and Tao. (b,d,f) Cross-compendium correlation matrices for Drews versus Tao, Drews versus Steele, and Steele versus Tao. Cross-compendium correlations use the 5,881 tumors represented in all three compendia after normalization of TCGA patient identifiers.

Cross-compendium concordance was selective rather than global (Figure 3b,d,f). Only 20 cross-framework pairs had Pearson correlations above 0.50, and six exceeded 0.60. Nine of these 20 pairs involved Steele and Tao, which is compatible with the shared use of copy number, loss-of-heterozygosity, and segment-size information in these frameworks. Strong relationships, including CN1 with Sig1, Sig2, and Sig5, CN9 with Sig10, and CN2 with CX6 (r = 0.70), occurred as isolated pairs rather than as broad matching blocks. These associations therefore identify selected shared axes of variation, but do not support one-to-one equivalence between signature systems. The corresponding correlation matrices restricted to the 12 high-activity signatures used for clustering are shown in Supplementary Figure 1.

The binary analysis addressed a different question: whether two signatures assigned the same patients to active or inactive states (Figure 4a, b). CN1 showed high agreement with Sig1 (approximately 0.76) and Sig2 (approximately 0.72), whereas CX1 showed moderate agreement with CN1 (approximately 0.58) and CN1 and CN2 showed little agreement (approximately 0.15). Because concordant inactivity contributed symmetrically to the metric, high agreement for sparse signatures could be driven by shared absence as well as shared activity. Patient-state agreement should therefore not be interpreted as evidence of coactivation or common aetiology. CX1 and CN1 illustrate this distinction: despite their moderate agreement, CX1 represents a chromosome- missegregation-associated pattern, whereas CN1 represents a copy-number-quiet diploid state. Together, the correlation and binary benchmarks reveal a limited number of shared patient-state patterns while rejecting broad interchangeability of the three compendia.

**Figure 4.**
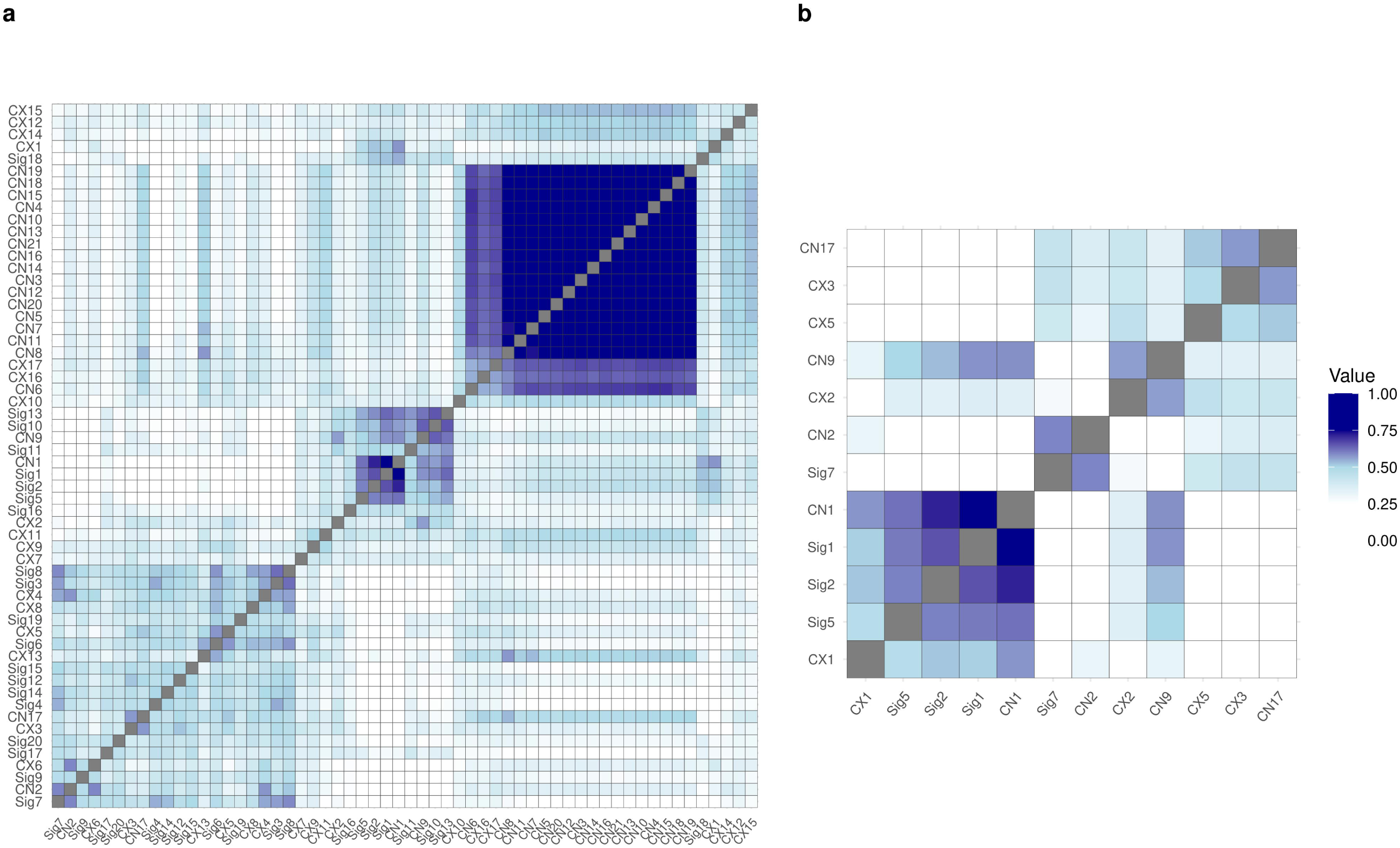
Binary agreement of copy number signature classifications. (a) Agreement matrix across all signatures after median-based binary classification. (b) Agreement matrix restricted to the high-activity signatures. Values show the Jaccard similarity of patient-level state assignments, A/(2N - A), across the 5,881 matched tumors. Here, A counts concordant active-active and inactive-inactive assignments and N is the number of matched tumors. Both states contribute symmetrically because activity and inactivity are informative classifications.

### Compendia generate distinct patient partitions

Model-based clustering of the high-activity signatures yielded four Drews clusters, four Steele clusters, and three Tao clusters (Figure 5a,d,g; Supplementary Figure 2). The retained cluster structures were driven by different signature combinations (Figure 5b,e,h), consistent with the distinct feature encodings of the three compendia.

**Figure 5.**
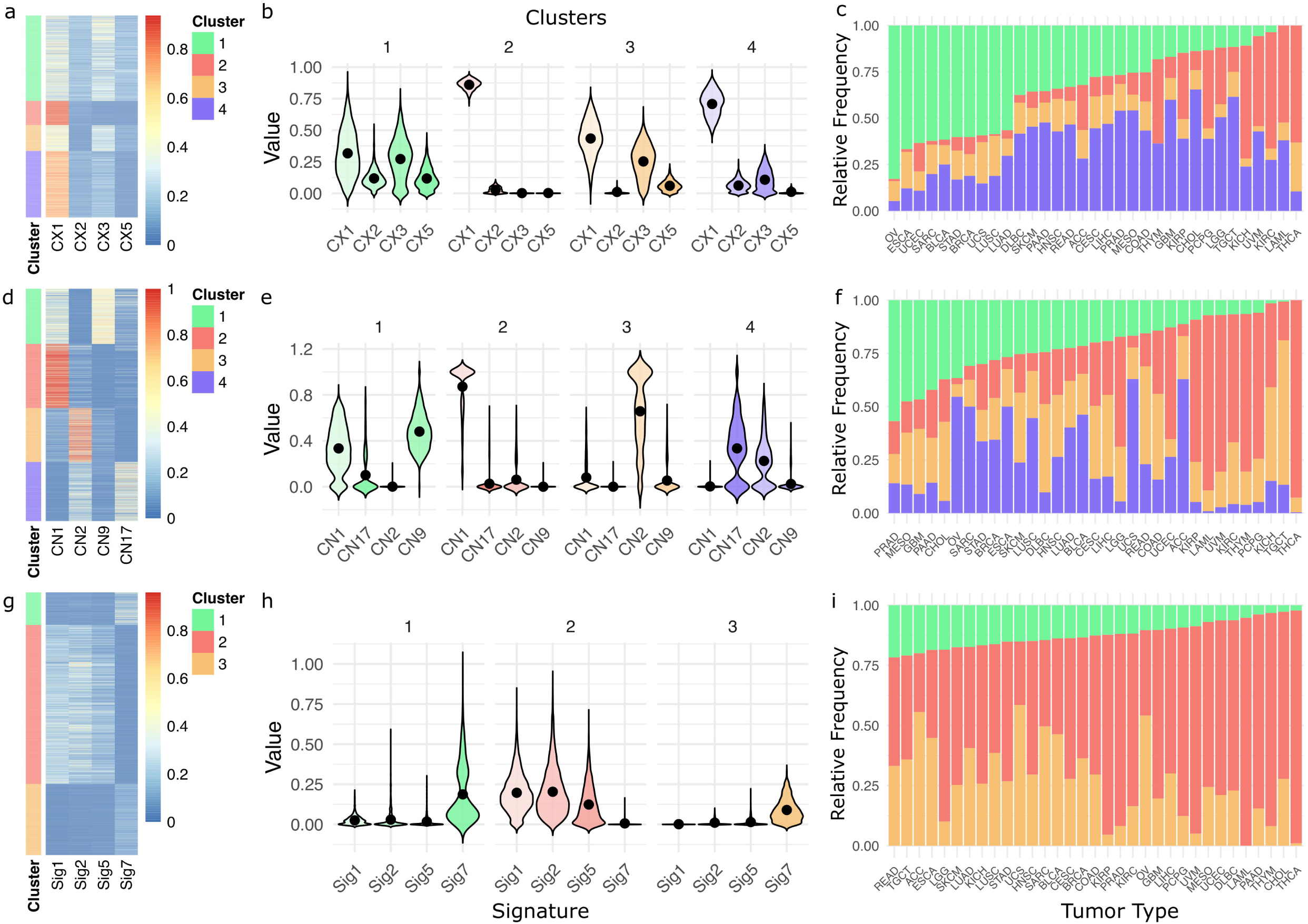
Compendium-specific clustering of high-activity copy number signatures. (a,d,g) Heatmaps of tumors ordered by model-based cluster for Drews, Steele, and Tao. (b,e,h) Violin plots of signature activity within each cluster. (c,f,i) Relative cluster composition within each cancer type. Four clusters were retained for Drews and Steele and three for Tao.

In Drews, Cluster 2 combined high CX1 activity with near-complete absence of CX2, CX3, and CX5. This pattern is best interpreted as CX1-dominant and depleted of the principal HRD-associated signatures, rather than as a generally stable genome. Cluster 4 had lower CX1 and partial CX2/CX3 activation, Cluster 3 combined moderate CX1 and CX3 activity, and Cluster 1 showed intermediate contributions from all four signatures. In Steele, Cluster 1 was dominated by CN9, a focal LOH and diploid-instability pattern; Cluster 2 was enriched for the diploid CN1 signature; Cluster 3 was dominated by the tetraploidy-associated CN2 signature; and Cluster 4 combined CN17, associated with HRD, with moderate CN2 activity. In Tao, Clusters 1 and 3 showed high and intermediate Sig7 activity, respectively, whereas Cluster 2 was Sig7 depleted and instead showed higher Sig1, Sig2, and Sig5 activities.

Cluster composition varied across cancer types but was not reducible to a one-to-one mapping between cluster and tissue of origin (Figure 5c,f,i). Some cancer types were predominantly represented in specific clusters, while others were distributed across multiple clusters. Cross-compendium comparison further showed limited overlap between identically numbered patient groups (Supplementary Figure 3). The largest concordance was observed between Steele and Tao, but only a minority of patients assigned to Cluster 2 were shared across all three frameworks. Thus, each compendium induced a distinct partition of the pan-cancer cohort.

### Outcome associations are directionally concordant but framework dependent

Kaplan-Meier analyses identified Cluster 2 as the group with the most favorable descriptive outcome profile in each compendium (Figure 6; Supplementary Figure 5). Because the distribution of cancer types differed among clusters, these curves alone could not establish a pan-cancer cluster association. We therefore used Cox models stratified by cancer type and adjusted for age and tumor purity.

**Figure 6.**
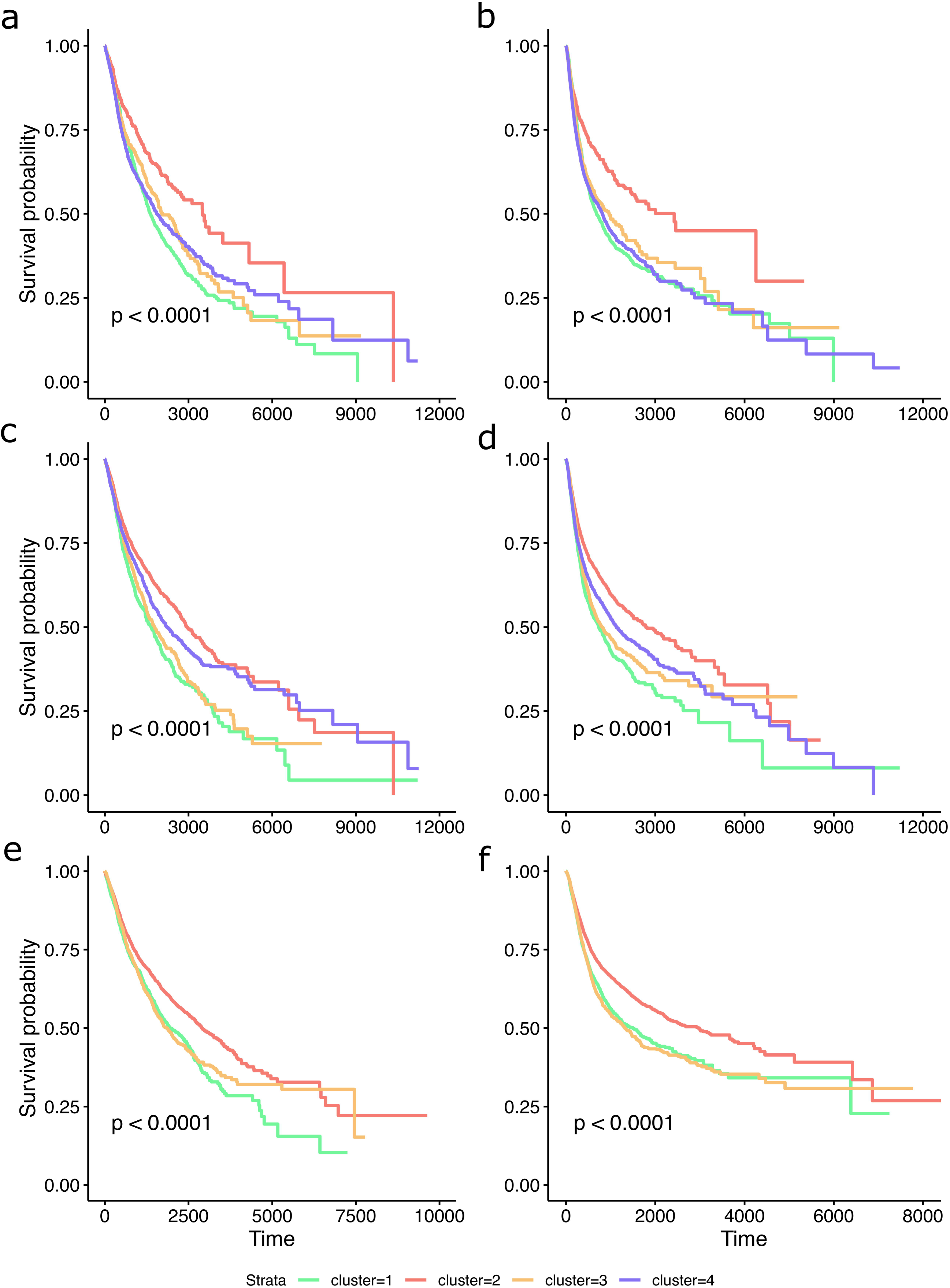
Survival distributions across signature-derived clusters. Kaplan-Meier curves show overall survival (a,c,e) and progression-free interval (b,d,f) for Drews (a,b), Steele (c,d), and Tao (e,f). Time is shown in days; corresponding numbers at risk are provided in Supplementary Figure 5.

The comparison of Cluster 2 with Cluster 1 showed a directionally favorable association in all six compendium- endpoint combinations, with varying effect size and statistical support (Figure 7). In Drews, Cluster 2 was associated with improved OS (HR = 0.64, 95% CI 0.54-0.77, Holm-adjusted P < 0.001) and PFI (HR = 0.68, 95% CI 0.58-0.80, Holm-adjusted P < 0.001). Steele showed smaller associations for OS (HR = 0.85, 95% CI 0.76-0.95, Holm-adjusted P = 0.034) and PFI (HR = 0.85, 95% CI 0.77-0.94, Holm-adjusted P = 0.021). In Tao, Cluster 2 was associated with improved PFI (HR = 0.87, 95% CI 0.79-0.95, Holm-adjusted P = 0.027), whereas the OS estimate was weaker and not statistically significant (HR = 0.92, 95% CI 0.84-1.01, Holm-adjusted P = 0.827).

**Figure 7.**
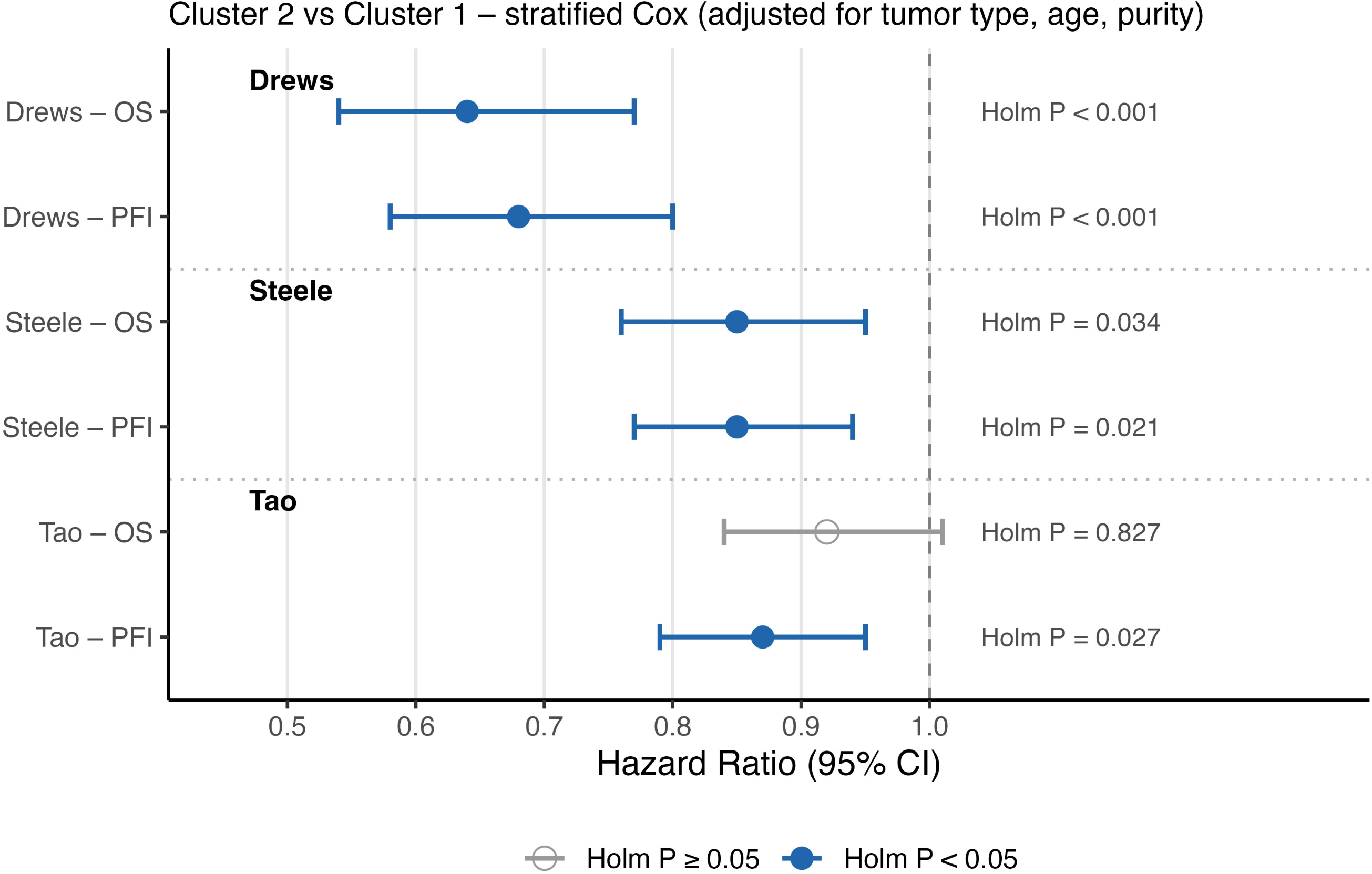
Cancer-type-stratified Cox benchmark of Cluster 2 versus Cluster 1. Points show hazard ratios and horizontal lines show 95% confidence intervals for overall survival and progression-free interval in the Drews, Steele, and Tao compendia. Models include age and tumor purity as continuous covariates and cancer type as a stratification variable. Two-sided Wald P values were adjusted across all 16 cluster coefficients from the six models using Holm’s method. Hazard ratios below 1 favor Cluster 2; filled blue points denote Holm-adjusted P < 0.05 and open gray points denote Holm- adjusted P ≥ 0.05.

Five of the six reported Cluster 2 versus Cluster 1 comparisons therefore retained statistical evidence of a cluster association after Holm correction across all 16 cluster coefficients. However, effect sizes differed between frameworks, and the favorable clusters contained substantially different patients. These findings support cluster-level clinical relevance as a secondary benchmark while arguing against treating any cluster assignment as a validated prognostic biomarker.

### Multi-omic predictability differs sharply by signature framework

We next evaluated whether the representations defined by each compendium were associated with molecular information beyond the copy number data used to derive them. This predictive analysis was restricted to TCGA sample-type code 01 (Primary Solid Tumor) and therefore excluded LAML, unlike the earlier signature comparison, clustering, and survival analyses. Continuous prediction of individual signature activities yielded heterogeneous and generally modest performance (Supplementary Figure 4). CX1, CX2, CX5, CN1, Sig2, and Sig5 were among the more predictable signatures, with the highest observed coefficient of determination reaching 0.65 for CX1 using ridge regression. Extreme gradient boosting and ridge regression were the strongest overall performers across these continuous tasks. We then tested whether the same multi-omic feature classes could predict the cluster assignments derived from each compendium.

Cluster-level prediction produced a clearer contrast among the three compendia. Using the same multi-omic feature classes and held-out evaluation design, the Cluster 1 versus Cluster 2 classifiers achieved test-set F1 scores of 0.93 for Drews, 0.80 for Steele, and 0.24 for Tao. Overall and cancer-type-specific test accuracies are shown in Figure 8a-c, while the corresponding test-set sample sizes are shown in Figure 8d. Accuracy should be interpreted cautiously because of cluster imbalance; the low Tao F1 score despite its comparatively high overall accuracy illustrates this limitation directly.

**Figure 8.**
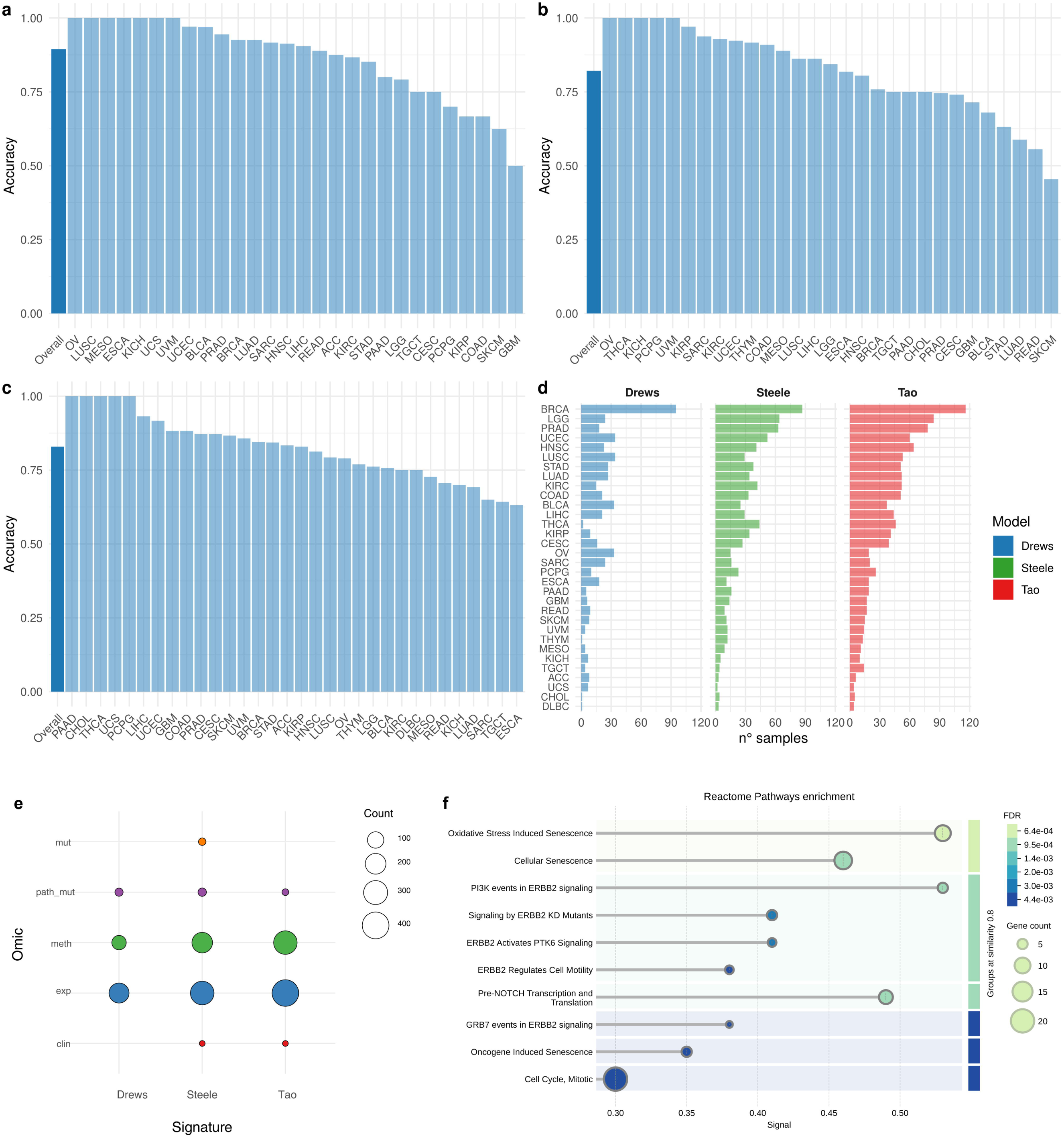
Framework-dependent multi-omic prediction, feature importance, and pathway enrichment. (a-c) Overall and cancer-type-specific test accuracy for Cluster 1 versus Cluster 2 classifiers trained for Drews, Steele, and Tao; cancer types with fewer than four test samples are omitted. (d) Test-set sample counts by cancer type and compendium. (e) Number of high-importance predictors from gene expression, DNA methylation, gene-level mutations, pathway-level mutations, and clinical variables. (f) Reactome terms significantly enriched among high-importance features from the Drews classifier; color denotes false discovery rate and point size denotes gene count.

Although cancer type was not included as a predictor, molecular profiles can encode tissue-of-origin information; therefore, these results do not establish complete independence from tumor lineage. Instead, they show that the Drews- and Steele-defined cluster boundaries are more readily recoverable from matched molecular profiles than the Tao-defined boundary under the same modeling framework.

### Predictive feature composition differs across frameworks

The number of variables required to account for 80% of cumulative model gain differed across frameworks: 300 for Drews, 576 for Steele, and 788 for Tao. The Drews classifier therefore concentrated 80% of its cumulative gain in a smaller predictor set, whereas predictive gain was distributed across more variables for Steele and Tao. Gene expression accounted for the largest number of retained variables in all three models, followed by DNA methylation; gene-level mutations, pathway-level mutations, and clinical variables were comparatively sparse (Figure 8e).

Given the prevalence of gene expression features in the retained predictor sets and their broad availability in public cohorts, we additionally evaluated classifiers restricted to gene expression. These models achieved test-set F1 scores of 0.94 for Drews, 0.78 for Steele, and 0.30 for Tao, preserving the same framework-dependent ranking observed with the full multi-omic classifiers.

Only the Drews retained feature set showed significant Reactome enrichment after false-discovery-rate correction (Figure 8f). Enriched terms included oxidative-stress-induced and oncogene-induced senescence, ERBB2-related signaling, pre-NOTCH transcription and translation, and mitotic cell-cycle processes. These pathways are compatible with DNA damage, replication stress, and chromosome-segregation phenotypes, but feature importance and enrichment remain associational and do not establish causal drivers of cluster membership.

Tumor purity was retained by the Steele classifier and was the highest-ranked predictor for Tao, whereas it was absent from the Drews 80%-gain feature set; the Drews source cohort had been restricted to higher-purity tumors. This difference may reflect both biological variation in microenvironment composition and technical dilution of tumor-specific molecular and copy number signals. Taken together, these results indicate that the recoverability of signature-defined clusters from non-copy-number molecular profiles depends on both the selected signature representation and cohort characteristics, rather than being a universal property of copy number signatures.

## Discussion

This study provides a matched-sample, multi-level benchmark of three established pan-cancer copy number signature compendia. Its central result is not the identification of a single superior signature system, but the demonstration that compendium choice changes the representation of genome instability, the partitioning of patients, the magnitude of outcome associations, and the extent to which those states can be reconstructed from other molecular data. The three frameworks should therefore be treated as complementary analytical models rather than interchangeable labels for the same underlying variable.

The sparse cross-compendium correlations are consistent with the way the signatures were constructed. Drews emphasizes CIN processes encoded through breakpoint and segment-pattern features, Steele incorporates allele-specific copy number and LOH to resolve broad genome states, and Tao adds local morphological context. Selected signatures converge when they capture closely related ploidy or LOH states, but broad one-to-one mapping is not expected. The binary analysis treats shared inactivity as informative patient-state agreement by design. For sparse signatures, high agreement may therefore be driven primarily by concordant inactive classification; this establishes consistency of state assignment, not necessarily coactivation or shared aetiology. The moderate agreement of CX1 and CN1 is a clear example, because a chromosome-missegregation-associated pattern and a copy-number-quiet diploid pattern are not mechanistic substitutes. More generally, copy number signatures summarize accumulated genomic states rather than directly measuring ongoing chromosome missegregation; comparative evaluations of CIN measures indicate that direct single-cell approaches are better suited to quantify active instability (23).

Cluster-level analyses reveal a second form of non-equivalence. Each compendium generated coherent patient groups, but the groups were composed of different tumors and were driven by different signature combinations. Even the three favorable-outcome Cluster 2 groups had limited shared membership. Their directionally concordant Cox estimates therefore do not demonstrate that the frameworks recover the same low-risk biological state. A more conservative interpretation is that distinct copy number representations can each summarize aspects of tumor genome organization associated with outcome. The effect was strongest for Drews, smaller for Steele, and incomplete for Tao, and all estimates arose from retrospective pan-cancer data. These clusters should consequently be regarded as comparative stratification outputs rather than clinically validated biomarkers.

The machine learning benchmark adds an orthogonal way to compare representations. Drews and Steele cluster states were recoverable from matched multi-omic profiles, whereas the Tao state was poorly recovered when F1 score was used to account for class imbalance. This difference persisted in expression-only models. One interpretation is that Drews and Steele produce cluster boundaries more closely aligned with broad transcriptional and epigenetic programs, whereas the Tao encoding retains finer copy number morphology that is less directly represented in these data layers. Differences in sample composition, purity, class balance, and target geometry may also contribute. The results therefore support predictability as a comparative property of a representation, not as evidence that expression or methylation can replace direct copy number measurement.

This distinction matters as copy number analysis moves toward new data types and clinical settings. Recent studies have inferred consensus signatures from targeted panel data (7), learned cancer-associated copy number patterns with attention-based models (8), and shown that systematic analysis of copy number gains can reveal therapeutic vulnerabilities (22). These advances expand the range of available representations, but they also increase the need for matched-sample comparison and explicit reporting of what a signature encodes. The present framework complements method-development studies by benchmarking established compendia at the signature, patient, clinical, and multi-omic levels within the same pan-cancer resource.

Several limitations define the scope of the conclusions. First, no independent cohort currently provides matched copy number signature activities for all three compendia together with uniformly processed expression, methylation, mutation, purity, and outcome data across many cancer types. The study therefore uses held-out test sets for predictive evaluation but lacks external validation of the complete benchmark. Second, all analyses are retrospective and based on public cohorts, and the pan-cancer survival models estimate a common cluster effect after allowing cancer-type-specific baseline hazards; they do not establish cancer-specific clinical utility. Third, clustering depends on the selected high-activity signatures and mixture-model solution, and alternative selection or clustering strategies may alter patient assignments. Fourth, the patient-state Jaccard gives equal weight to concordant active and inactive classifications and therefore measures overall state-assignment agreement rather than positive-state overlap alone. Fifth, the machine learning comparison used a single cancer-type-stratified train-test partition, and residual tissue-of-origin information can remain in molecular profiles even when the cancer-type label is excluded. Finally, feature importance and pathway enrichment are descriptive and do not establish causal mechanisms.

Within these constraints, the benchmark establishes a practical conclusion: current copy number signature compendia preserve different, partially overlapping views of tumor genome instability. In practice, the relevant choice is not which compendium is universally superior, but whether the intended analysis prioritizes process-oriented CIN patterns (Drews), allele-specific genome states (Steele), or local copy number morphology (Tao). Explicitly benchmarking concordance, patient partitioning, clinical associations, and molecular predictability provides a more defensible basis for choosing among them than assuming equivalence. Independent, uniformly profiled cohorts will be required to determine which framework-specific states generalize and whether any are suitable for prospective biomarker development.

## Data Availability

The copy number signature activity matrices were obtained from the repositories accompanying the original Drews, Steele, and Tao publications (3–5). Gene expression and DNA methylation data were obtained through curatedTCGAData; somatic mutation data were obtained from the public MC3 mutation annotation file; clinical outcomes were obtained from the pan-cancer clinical resource of Liu et al. (12); and tissue source site codes were obtained from the Genomic Data Commons.

All analysis code is openly available at https://github.com/marcorotanegroni/CNS_ML. A versioned archive of the code and processed multi-omic datasets needed to reproduce the analyses and retrain the predictive models is available from Zenodo at https://doi.org/10.5281/zenodo.20617051.

## Supplementary Data

Supplementary Data are available Online.

## Acknowledgements

The results shown here are in whole or in part based upon data generated by the TCGA Research Network: https://www.cancer.gov/tcga.

## Author Contributions

Marco Rota Negroni: Data curation, Formal analysis, Methodology, Software, Visualization, Writing - original draft. Ilaria Billato: Methodology, Writing - review and editing. Chiara Romualdi: Conceptualization, Funding acquisition, Supervision, Writing - review and editing. All authors read and approved the submitted version.

## Funding

This work was supported by the Italian Association for Cancer Research [IG 29071 to C.R.]; and by the European Union - NextGenerationEU through the Italian Ministry of University and Research, National Recovery and Resilience Plan, Mission 4 Component 2, Action 1.4, National Center for HPC, Big Data and Quantum Computing [CN00000013, Spoke 8].

## Conflict of Interest

The authors declare no competing interests.

## Notes

https://github.com/marcorotanegroni/CNS_ML

### Competing Interest Statement

The authors have declared no competing interest.

### Summary of Updates

No new data were generated and no new analyses were run. The title, abstract, introduction and discussion were rewritten so that the study is presented as a matched sample benchmark of three published copy number signature compendia, with compendium choice framed as an analytical decision rather than an interchangeable preprocessing step. The P values for the 16 cluster coefficients estimated across the six compendium and endpoint models are now adjusted as a single family using the Holm method, so the reported P values differ from version 1 although the point estimates do not. Several claims were qualified rather than changed, additional references were cited, and the limitations section was expanded.

