## Supplementary Data for "Pan-cancer benchmarking reveals complementary copy number signatures with distinct multi-omic predictability"

### Supplementary Table 1

Number and percentage of samples with available signature estimates. Each row corresponds to a different cancer type, while the three main columns refer to the Drews, Steele, and Tao signature models. For TCGA cancer type codes, see the reference here: https://gdc.cancer.gov/resources-tcga-users/tcga-code-tables/tcga-study-abbreviations

| | Drews | | Steele | | Tao | |
|---|---|---|---|---|---|---|
| Tumor type | n° samples | % Total | n° samples | % Total | n° samples | % Total |
| ACC | 78 | 1.23 | 89 | 0.92 | 90 | 0.87 |
| BLCA | 260 | 4.10 | 372 | 3.84 | 391 | 3.77 |
| BRCA | 730 | 11.52 | 1020 | 10.52 | 1066 | 10.28 |
| CESC | 220 | 3.47 | 292 | 3.01 | 299 | 2.88 |
| CHOL | 29 | 0.46 | 35 | 0.36 | 36 | 0.35 |
| COAD | 293 | 4.63 | 413 | 4.26 | 452 | 4.36 |
| DLBC | 24 | 0.38 | 41 | 0.42 | 48 | 0.46 |
| ESCA | 132 | 2.08 | 176 | 1.81 | 183 | 1.76 |
| GBM | 427 | 6.74 | 500 | 5.16 | 504 | 4.86 |
| HNSC | 308 | 4.86 | 492 | 5.07 | 517 | 4.99 |
| KICH | 46 | 0.73 | 66 | 0.68 | 66 | 0.64 |
| KIRC | 167 | 2.64 | 467 | 4.81 | 517 | 4.99 |
| KIRP | 95 | 1.50 | 229 | 2.36 | 284 | 2.74 |
| LAML | 21 | 0.33 | 113 | 1.17 | 190 | 1.83 |
| LGG | 285 | 4.50 | 472 | 4.87 | 508 | 4.90 |
| LIHC | 283 | 4.47 | 344 | 3.55 | 369 | 3.56 |
| LUAD | 239 | 3.77 | 477 | 4.92 | 507 | 4.89 |
| LUSC | 302 | 4.77 | 475 | 4.90 | 493 | 4.75 |
| MESO | 59 | 0.93 | 82 | 0.85 | 86 | 0.83 |
| OV | 545 | 8.60 | 558 | 5.75 | 558 | 5.38 |
| PAAD | 65 | 1.03 | 133 | 1.37 | 181 | 1.75 |
| PCPG | 90 | 1.42 | 153 | 1.58 | 162 | 1.56 |
| PRAD | 285 | 4.50 | 435 | 4.48 | 486 | 4.69 |
| READ | 133 | 2.10 | 148 | 1.53 | 166 | 1.60 |
| SARC | 196 | 3.09 | 230 | 2.37 | 248 | 2.39 |
| SKCM | 348 | 5.49 | 442 | 4.56 | 103 | 0.99 |
| STAD | 231 | 3.65 | 427 | 4.40 | 436 | 4.20 |
| TGCT | 104 | 1.64 | 143 | 1.47 | 134 | 1.29 |
| THCA | 19 | 0.30 | 248 | 2.56 | 498 | 4.80 |
| THYM | 11 | 0.17 | 77 | 0.79 | 123 | 1.19 |
| UCEC | 221 | 3.49 | 424 | 4.37 | 536 | 5.17 |
| UCS | 54 | 0.85 | 54 | 0.56 | 53 | 0.51 |
| UVM | 35 | 0.55 | 72 | 0.74 | 80 | 0.77 |

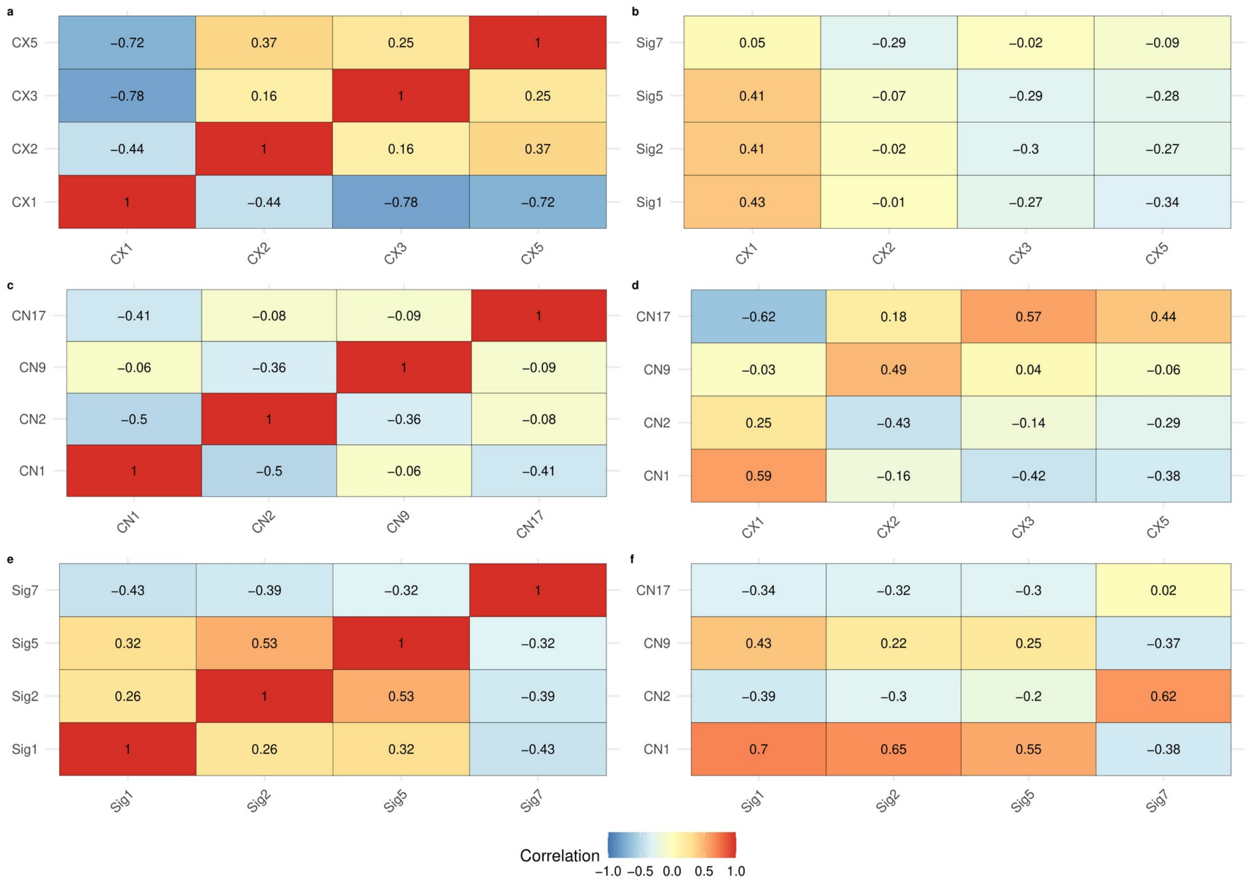


**Supplementary Figure 1. Within- and cross-compendium Pearson correlation analysis of high-activity copy number signatures.** (a, c, e) Within-compendium Pearson correlation matrices for the high-activity signatures defined in the Drews, Steele, and Tao compendia, respectively. (b, d, f) Cross-compendium Pearson correlation matrices comparing the high-activity signature profiles between Drews and Tao (b), Drews and Steele (d), and Steele and Tao (f).

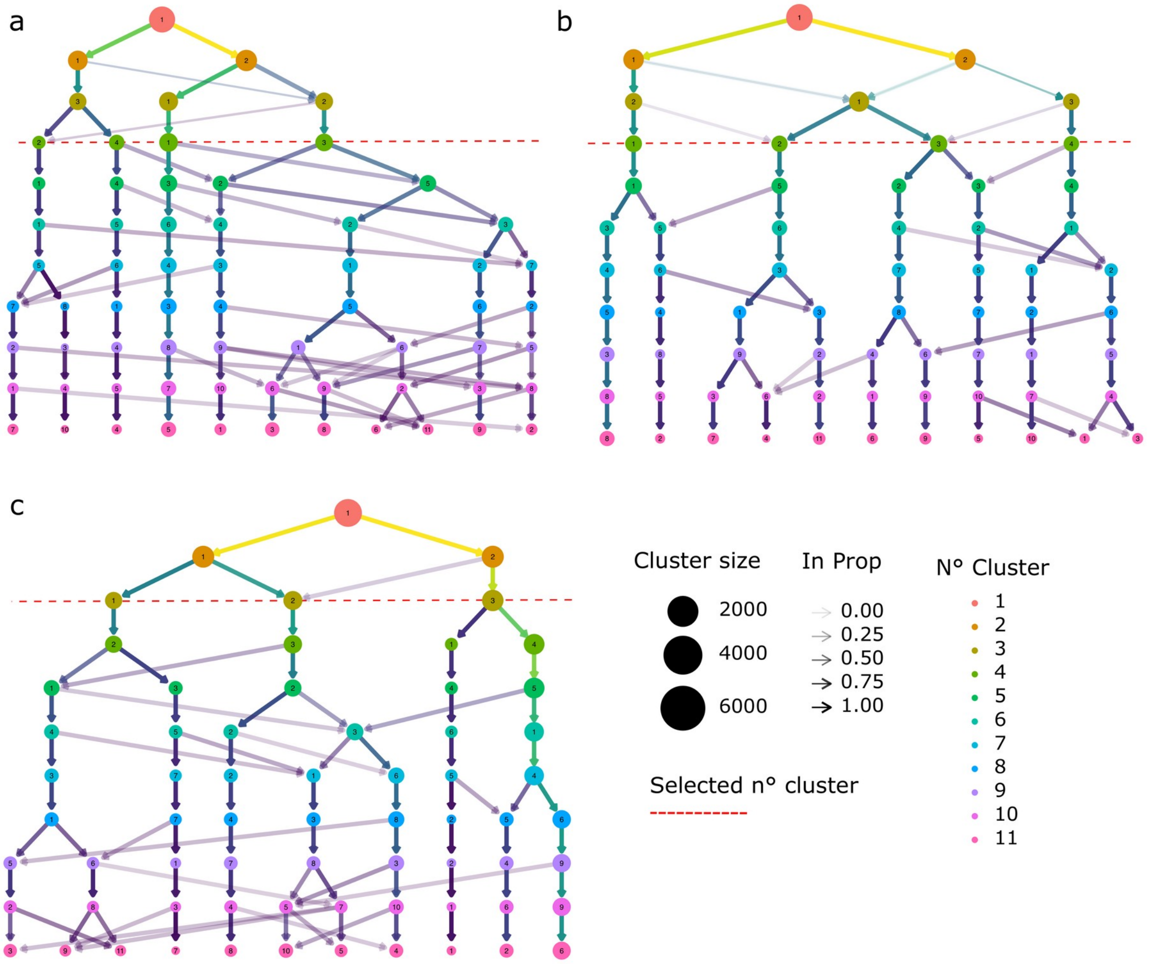


**Supplementary Figure 2. Clustering trees used to inspect candidate patient partitions.** Nodes are arranged in layers corresponding to exploratory k-means solutions with increasing k, and edges show how samples move between resolutions. Edge width and color are proportional to the in-proportion metric, defined as the number of samples moving from one cluster to another as k increases. Dashed red lines indicate the selected cluster numbers for Drews (a), Steele (b), and Tao (c). These exploratory trees guided the candidate range; final classifications in the main analysis were obtained by Gaussian finite mixture modeling.

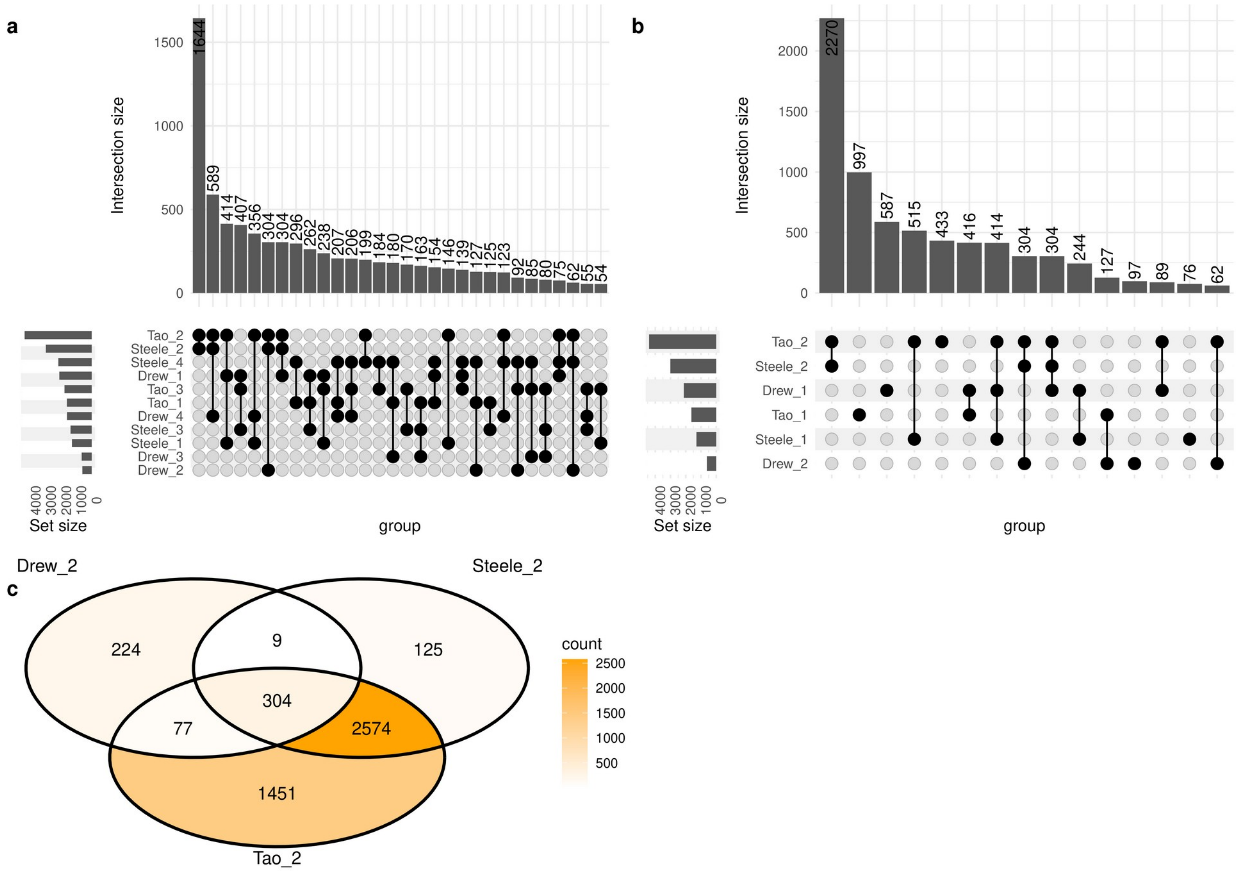


**Supplementary Figure 3. Overlap of signature-based clusters across studies.** (a) UpSet plot illustrating sample overlap among clusters defined in the three datasets. (b) Enlarged view of sample overlap restricted to Clusters 1 and 2, which showed less and more favorable descriptive survival profiles, respectively. (c) Venn diagram showing the intersection of Cluster 2 samples across studies.

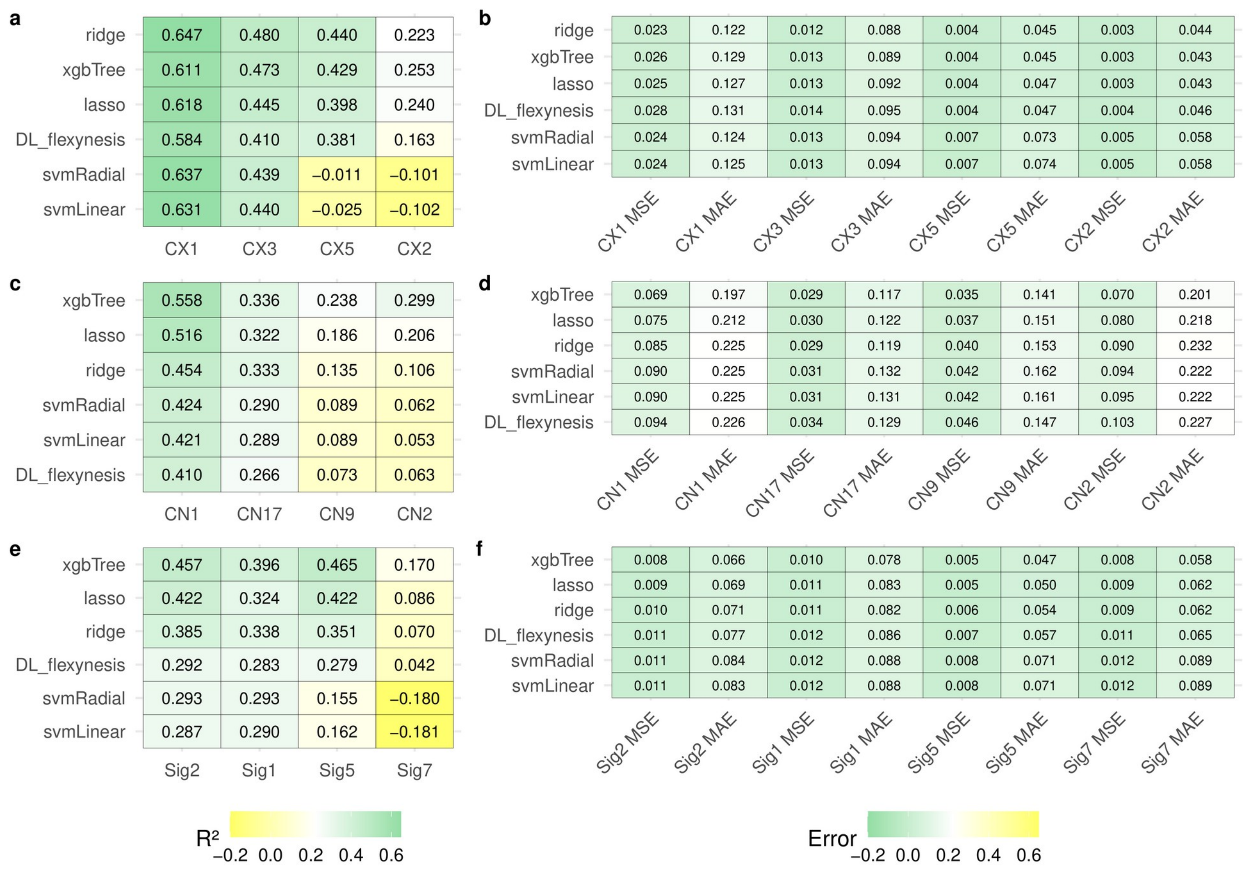


**Supplementary Figure 4. Comparative performance of regression models for continuous signature prediction.** (a,c,e) Coefficients of determination ($R^2$) for Ridge, Lasso, support vector machines with linear and radial kernels, extreme gradient boosting, and deep learning models applied to selected Drews (a), Steele (c), and Tao (e) signatures. (b,d,f) Corresponding mean absolute error and mean squared error values for Drews (b), Steele (d), and Tao (f). Models are ranked from top to bottom by mean coefficient of determination or minimum error. Red intensity indicates higher $R^2$ or lower error.

a

Number at risk

| Strata | 0 | 3000 | 6000 | 9000 | 12000 |
|---|---|---|---|---|---|
| cluster=1 | 2629 | 137 | 12 | 1 | 0 |
| cluster=2 | 740 | 56 | 5 | 1 | 0 |
| cluster=3 | 797 | 49 | 7 | 1 | 0 |
| cluster=4 | 2041 | 110 | 16 | 2 | 0 |

b

Number at risk

| Strata | 0 | 3000 | 6000 | 9000 | 12000 |
|---|---|---|---|---|---|
| cluster=1 | 2629 | 99 | 8 | 0 | 0 |
| cluster=2 | 730 | 46 | 4 | 0 | 0 |
| cluster=3 | 796 | 30 | 4 | 1 | 0 |
| cluster=4 | 2032 | 65 | 8 | 2 | 0 |

c

| Strata | 0 | 3000 | 6000 | 9000 | 12000 |
|---|---|---|---|---|---|
| cluster=1 | 1617 | 72 | 5 | 1 | 0 |
| cluster=2 | 3628 | 194 | 14 | 2 | 0 |
| cluster=3 | 1647 | 87 | 6 | 0 | 0 |
| cluster=4 | 2609 | 167 | 25 | 4 | 0 |

d

| Strata | 0 | 3000 | 6000 | 9000 | 12000 |
|---|---|---|---|---|---|
| cluster=1 | 1610 | 47 | 3 | 1 | 0 |
| cluster=2 | 3536 | 151 | 7 | 0 | 0 |
| cluster=3 | 1646 | 56 | 6 | 0 | 0 |
| cluster=4 | 2599 | 118 | 16 | 2 | 0 |

e

| Strata | 0 | 2500 | 5000 | 7500 | 10000 |
|---|---|---|---|---|---|
| cluster=1 | 2110 | 181 | 22 | 1 | 0 |
| cluster=2 | 6184 | 493 | 35 | 5 | 0 |
| cluster=3 | 1939 | 156 | 7 | 0 | 0 |

Time in days

f

| Strata | 0 | 2000 | 4000 | 6000 | 8000 |
|---|---|---|---|---|---|
| cluster=1 | 1929 | 187 | 17 | 3 | 0 |
| cluster=2 | 6011 | 630 | 69 | 8 | 3 |
| cluster=3 | 2110 | 200 | 34 | 11 | 0 |

Time in days

**Supplementary Figure 5. Kaplan-Meier survival risk tables.** Risk tables corresponding to overall survival (a,c,e) and progression-free interval (b,d,f) for the signature-derived clusters in Drews (a,b), Steele (c,d), and Tao (e,f), as shown in Figure 6 of the main text.

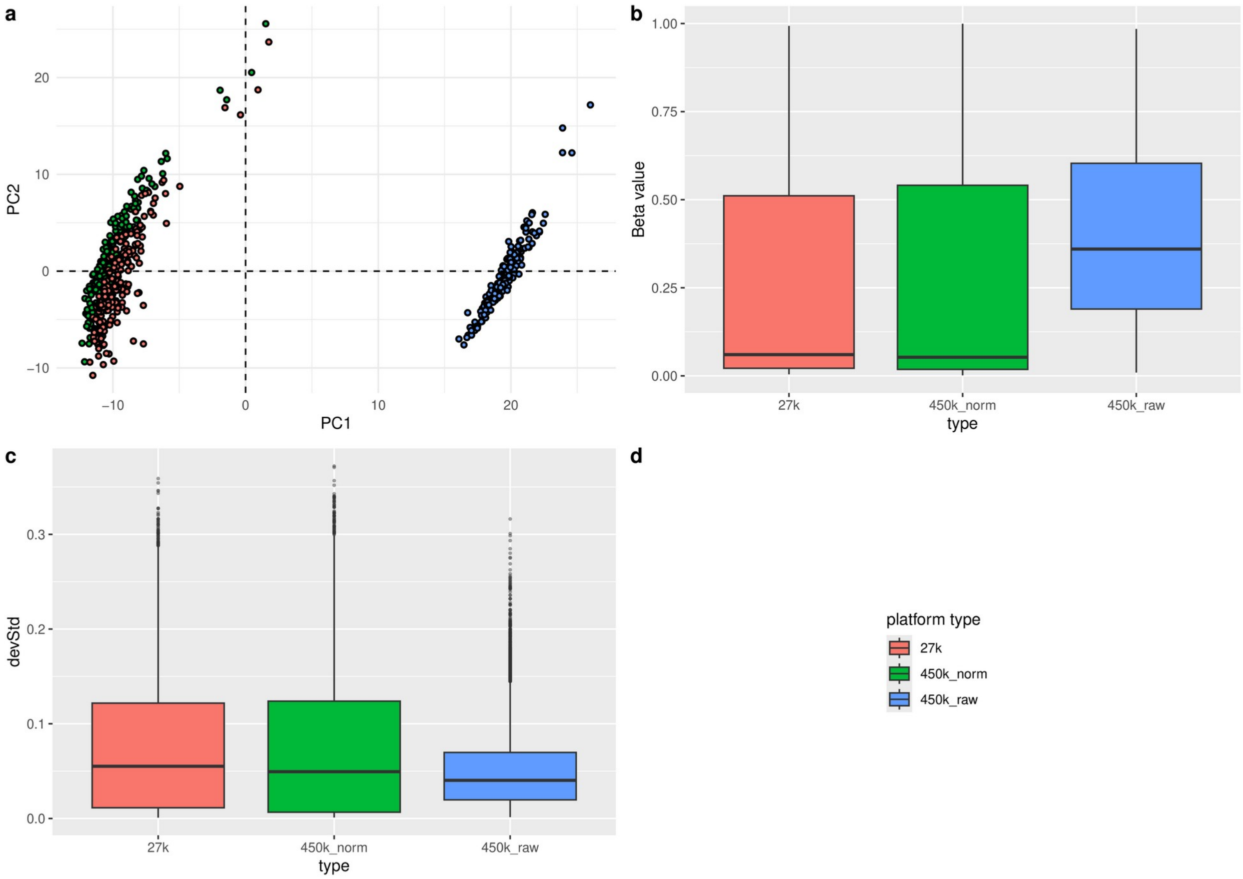


**Supplementary Figure 6. Harmonization of multi-platform DNA methylation data.** (a) Principal-component projection of acute myeloid leukemia methylation profiles from the 27k array (red), 450k array (blue), and beta-mixture-quantile-normalized 450k data (green), showing improved cross-platform alignment after normalization. (b) Beta-value distributions before and after normalization. (c) Standard-deviation profiles used to assess variance alignment across platforms after normalization.
